# Analgesic effects of neurotensin agonists in a rat bone cancer pain model

**DOI:** 10.1101/089169

**Authors:** Louis Doré-Savard, Pascal Tétreault, Mélisange Roux, Marylie Martel, Myriam Lemire, Karine Belleville, Nicolas Beaudet, Philippe Sarret

## Abstract

Bone metastases are a source of intractable pain, resistant to conventional opioid and non-opioid analgesics. The neurotensin system represents a potential pathway toward bone cancer pain (BCP) relieve via the inhibition of its receptors NTS1 and NTS2. Capitalizing on our recent results using neurotensin analogs in inflammatory and neuropathic pain models, we here show, for the first time, a potential role for neurotensin receptors agonists in the treatment of BCP. The novel non-selective agonist JMV-2009 (300 µg/kg) reversed mechanical allodynia in our rodent BCP model at both early and late stages of the disease. The NTS2-selective agonist JMV-431 (90 µg/kg), in addition to anti-allodynia, also had an effect on weight bearing deficits. In parallel, we tested proven analgesics from several classes to put the effect of neurotensin analogs in perspective and found that morphine (3 mg/kg), tramadol (15 mg/kg) and amitriptyline (10 mg/kg) had mild effects on BCP while the cannabinoid nabilone (1 mg/kg) significantly reversed both allodynia and weight bearing deficits. Taken together, our results affirm the potential of the modulation of the neurotensin system for the development of new analgesics for the treatment of bone cancer pain.

**Summary statement:** *Opioid and non-opioid analgesics were used to alleviate pain in a new rodent model of bone metastasis. The effect of each compound depended on the context and the evaluation method*.

## Introduction

Bone metastases are one of the most debilitating complications consecutive to primary soft organ cancers(Mercadante and Fulfaro, 2007). Pain frequently arises from these secondary tumors and progresses in intensity throughout the course of the disease. Indeed, advanced bone malignant pain is poorly managed, especially because of spontaneous breakthrough pain episodes(Mercadante et al., 2002). Non-steroidal anti-inflammatory drugs (NSAIDS) and opioids remain the gold standard for malignant skeletal pain(Irvin et al., 2011). However, doses of morphine required for providing a significant relief are reported to be as much as 10 times higher than for inflammatory syndromes(Luger et al., 2002). Moreover, the side effects associated to the opioid medication often lead to low compliance or treatment abortion.

With the recent findings that bone cancer pain could be associated with extended peripheral nerve damage and plasticity(Bloom et al., 2011; Peters et al., 2005), alternative opioid-independent approaches have been tested to address the neuropathic-like characteristics of the disease(Irvin et al., 2011). Among those, cannabinoids and multivalent compounds such as tramadol and tricyclic antidepressants have encountered various levels of success in both animal and human studies. But so far, they have mostly been considered as adjuvants for opioid regimen(Lussier et al., 2004). One of the emerging avenues in chronic pain, especially neuropathic, is the neurotensin (NT) system(Dobner, 2006). Our recent studies in several pain models uncovered an important role for two of the NT receptors, NTS1 and NTS2 in spinal and supraspinal pain inhibition. NTS1 stimulation efficiently reversed heat hyperalgesia and mechanical allodynia in a chronic neuropathic injury model(Guillemette et al., 2012). Moreover, NTS2-selective agonist JMV-431 produced a robust dose-dependant analgesia in the same model(Tétreault et al., 2013) and our group previously reported that selective NTS1 and NTS2 agonists both alleviated tonic inflammatory pain in rats(Demeule et al., 2014; Roussy et al., 2008; 2009).

Based on the analgesic actions of NT and considering the fact that BCP presents with neurochemical modifications at the spinal level consistent with inflammatory and neuropathic pain(Honore et al., 2000), we hypothesize that NT analogs should be efficient in decreasing BCP. In this study, we verified the spinal efficacy of a NTS1/NTS2 mixed analog and an NTS2-specific analog (JVM-2009 and JMV-431 respectively). In parallel, we validated the action of four clinically recommended drugs in treating different components of cancer pain in our specific model of rodent bone cancer pain. Neurotensin analogs along morphine, tramadol, amitriptyline and nabilone were evaluated for their efficiency to reverse mechanical allodynia and weight bearing deficits at early and late stages of the model.

## Materials and Methods

### Cell culture, animals and surgical procedure

Mammary rat metastasis tumor (MRMT-1) cells (carcinoma) were kindly provided by the Cell Resource Center for Biomedical Research Institute of Development, Aging and Cancer (Tohoku University 4-1, Seiryo, Aoba-ku, Sendai, Japan) and they were cultured and prepared according to our previous study (Doré-Savard et al., 2010). The cells were diluted to achieve a final concentration of 30,000 cells in 20 µL and maintained on ice prior to surgery.

Adult female Sprague-Dawley rats (200–225 g; Charles River Laboratories, St.-Constant, Quebec, Canada) were maintained on a 12 hrs light/dark cycle with access to food and water ad libitum. Animal-related procedures were approved by the Ethical Committee for Animal Care and Experimentation of the Université de Sherbrooke and conducted according to the regulations of the Canadian Council on Animal Care (CCAC). The rats were acclimatized to the animal facility for 4 days and to the manipulations and devices prior to the behavioral studies for 2 days.

The surgical procedure was performed as we previously described(Doré-Savard et al., 2010). Briefly, it consisted in the intrafemoral injection of 30 000 MRMT-1 cells without any damage to the joint. The sham animals received the complete surgical procedure except for the implantation of mammary cells, which was replaced by vehicle injection (20 µl HBSS). The rats were housed individually for 24 hours to allow for recovery.

### Imaging

Imaging procedures were described previously (Doré-Savard et al., 2010; 2012). In brief, MRI studies were conducted at the Centre d’imagerie moléculaire de Sherbrooke (CIMS) with a 210 mm small-animal 7T scanner (Agilent Technologies Inc., Palo Alto, CA, USA) and a 63 mm volume RF coil. The MRI protocol included the acquisition of axial (sagittal) pre-contrast and 10 min post-contrast T1-weighted images and a 500 ml bolus of the contrast agent (Gd-DPTA, Magnevist, Berlex) was injected via the tail vein.

### Drugs and injections

The animals were randomly assigned to treatment or vehicle groups following baseline behavioral measurements. The experimenter was blinded to treatment. The efficiency of acute treatment using proven analgesics or NT receptors agonists was evaluated in bone tumor–implanted rats on post-operative days 14 and 18. As previously determined(Doré-Savard et al., 2010), the day 14 was used to represent the early stage of BCP progression while the day 18 was identified as the late stage. Table 1 presents the six drugs used in our study along with the route of administration and references that helped us determining the appropriate dose, when applicable.

**Table 1.** List of drugs used in the present study.

| Drug | Route of administration | Dose | Elapsed time before tests | References | Source |
| --- | --- | --- | --- | --- | --- |
| Morphine sulfate | s.c. | 3 mg/kg | 30 minutes | (Medhurst et al., 2002) | Sabex Inc., Boucherville, QC, Canada |
| Amitriptyline | i.p. | 10 mg/kg | 120 minutes | (Mouedden and Meert, 2007) | Sigma-Aldrich, St-Louis, MO, USA |
| Tramadol | i.p. | 15 mg/kg | 30 minutes | (Cannon et al., 2010) | Sigma-Aldrich, St-Louis, MO, USA |
| Nabilone | *Per os* | 1 mg/kg | 60 minutes | (Guindon, 2012) and dose-response curve in acute pain model (not shown) | Ranbaxy Pharmaceuticals Canada, Mississauga, ON, Canada |
| JMV-431 | i.t. | 90 µg/kg | 30 minutes | (Doré-Savard et al., 2008; Roussy et al., 2009) | Kindly provided by Jean Martinez, Montpellier, France. |
| JMV-2009 | i.t. | 300 µg/kg | 30 minutes | Unpublished experiments in neuropathic and inflammatory pain (not shown) | Kindly provided by Jean Martinez, Montpellier, France |

All drugs were dissolved in physiological saline, except for Nabilone (Simple syrup B.P.: sucrose 0.88 g/ml and methylparaben 1.1 mg/ml). Volumes of injection were; 25 µl for the i.t. route, 1 ml/kg for the s.c. and i.p. routes and 5 ml/kg for the p.o. route.

### Behavioral evaluation of bone cancer pain

The animals were tested on days 0, 7, 11, 14 and 18 after cell implantation. For von Frey testing, the rats were placed in enclosures on an elevated wire mesh floor, and mechanical allodynia was assessed using a dynamic plantar aesthesiometer (Ugo Basile, Stoelting, IL, USA). A metal probe (0.5 mm diameter) was directed against the hind paw pad, and an upward force was exerted (3.33 g/second). The force required to elicit a withdrawal response was measured in grams and automatically registered when the paw was withdrawn or the preset cut-off was reached (50 g). Five values were taken alternately on both the ipsilateral and contralateral hind paws at intervals of 15 s. The rats were acclimatized to the enclosures for 2 days prior to testing.

The dynamic weight bearing (DWB) assessment was performed according to the procedure described previously(Tétreault et al., 2011). In brief, the dynamic weight bearing (DWB) device (Bioseb, Boulogne, France) consisted of a Plexiglas enclosure (22 × 22 × 30 cm) with a floor sensor composed of 44 × 44 captors (10.89 mm2 per captor). A digital camera was placed beside the cage and was pointed towards the enclosure. The rats were allowed to move freely within the apparatus for 5 min. while the pressure data and live recording were transmitted to a laptop computer via a USB interface. The raw pressure and visual data were colligated with the latest available DWB software (Bioseb). A zone was considered valid when the following parameters were detected: ≥ 4 g on 1 captor with a minimum of 2 adjacent captors recording ≥ 1 g. For each time segment in which the weight distribution was stable for more than 0.5 sec, the zones that met the minimal criteria were then validated and assigned as either right or left hind paw or front paw by an observer according to the video and the scaled map of the activated captors. Finally, a mean value for the weight on every limb was calculated for the whole testing period based on the length of time of each validated segment. The animals were not acclimatized to the enclosure before the initial testing period to maximize the exploration behaviors.

The percentage of anti-allodynia was calculated with the use of the following equation: % anti-allodynia = 100 × [(BCP treated rat – BCP vehicle-treated rat)/(Baseline - BCP vehicle-treated rat)]. From the latter formula, 0% represents no anti allodynic effect of the compound, while 100% corresponds to a complete relief of mechanical hypersensitivity. We used the same formula to calculate the % of rehabilitation and weight recovery.

### Quantitative real-time PCR

Lumbar DRG L1, L2 and L3 and corresponding spinal cord bulge located in the thoracic region were harvested at specific time point (sham and cancer at 18 days post-surgery) and processed as described earlier(Tétreault et al., 2013) to quantify the mRNA expression of NT, NTS1 and NTS2.

### Statistical analysis

An unpaired, non-parametric t-test (Mann-Whitney) was used to compare drug-treated to vehicle-treated animals in terms of % of anti-allodynia, and weight recovery. The same test was used for qPCR experiments. p < 0.05 was considered to be significantly different.

## Results

### Characterization of early and late stages in a bone cancer pain model

Tumor growth and pain development were monitored over time using several medical imaging techniques (Figure 1A) and behavioral testing (Figure 1B-C) respectively. Magnetic resonance imaging (MRI) provided a high-resolution visualization of tumor growth from its early settlement in bone marrow and cortex (Day 14) to the late extensive invasion of the bone and surrounding tissues (Day 18). On ^18^F-NaF positron emission tomography (PET), the radiotracer accumulates in metabolically active bone regions such as the metaphyses of the tibia and the femur, showing basal resorption/formation equilibrium in the sane bones. On Day 14 after tumor cells implantation, the tumor has invaded the bone marrow and modified trabecular and cortical bone. At this stage, ^18^F-NaF PET showed an increased uptake in the metaphysis and distal diaphysis. On co-registered images, it was obvious that active turnover was observed in bone subregions where the tumor was in contact with bone matrix. Active bone resorption and compensative sclerotic activity have been confirmed in these regions using histological observations(Doré-Savard et al., 2010; 2012). Bone modifications at the trabecular (not shown) and cortical levels are already observed on µCT. On day 18, contrast enhancement on MRI and the extent of tumor progression beyond bone were maximal. On PET images, ^18^F-NaF uptake was strongly decreased in the epiphysis and metaphysis while it was increased at the medial diaphysis level, where bone resorption is maximized. The striking degradation of bone matrix observed on µCT is in accordance with the important decreased ^18^F-NaF uptake.

**Figure 1.**
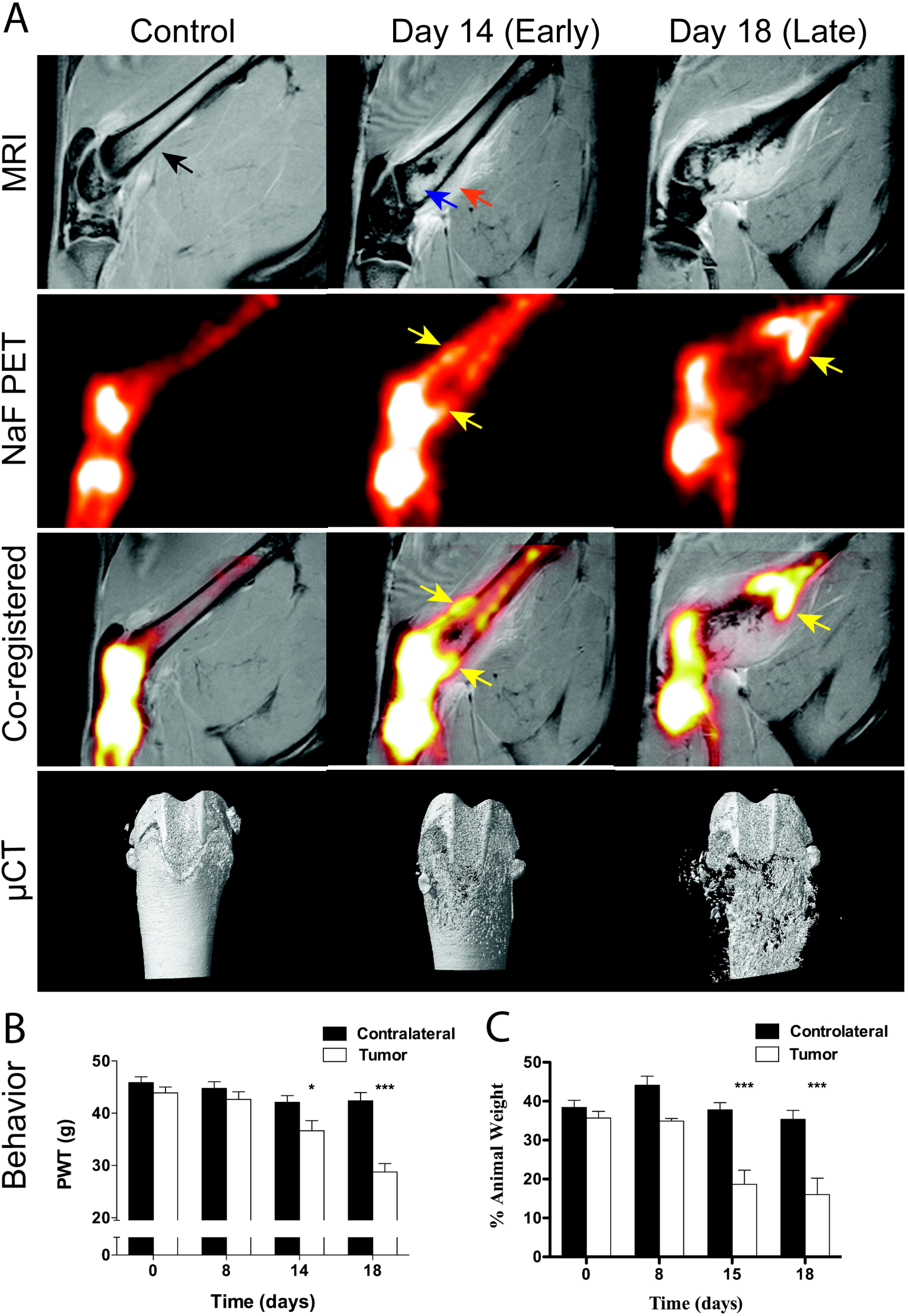
Basic characteristics of the BCP model. A) MRI shows progressive tumor growth over the course of the experiment. In control femurs (F), trabecular bone and cortical bone lines (black arrow) show smooth appearance and even contrast for bone marrow. On day 14, the tumor partially invaded the medullar cavity, as shown by increased intensity on the distal femur (blue arrow). The tumor also emerged from the bone (orange arrow). On day 18, extensive bone degradation and contrast enhancement is observed and tumor size compromises bone integrity. Areas of active bone resorption and compensative osteoblastic activity are shown on NaF PET images (yellow arrows) and co-registered MR/PET. The high level of bone resorption is also observed on µCT images were altered bone cortex is observed on day 14 and pathological fractures and isolated fragments are shown on day 18. B) Mechanical allodynia develops progressively from day 14. At day 18, Mechanical hypersensitivity reaches its maximal level. C) Concurrently, the animal shows weight bearing deficits on day 14 that persist over time until day 18.

The relationship between tumor volume, bone resorption and pain intensity has recently been described(Doré-Savard et al., 2012). Here we only show the timely progression of pain levels at relevant time points for this study using two behavioral assays; automated von Frey (Figure 1B) and Dynamic Weight Bearing (DWB) (Figure 1C). Mechanical allodynia was first detected on day 14 when the PWT was significantly decreased compared to the contralateral paw. After that early pain development, the PWT was again markedly decreased on day 18, considered as a late stage of tumor progression. On the DWB test, we also observed a significant decrease in the amount of weight bore by the tumor-affected paw, compared to the contralateral side. This postural deficit remained consistent between the early and the late stages of tumor growth. As it was mentioned before, analgesic drugs were administered at two different time points, being the early development of pain (day 14) and the late, more intense pain stages (day 18).

### Anti-allodynic effects of neurotensin agonists and other analgesics

We first evaluated the ability of neurotensinergic agonists to reverse mechanical allodynia using the von Frey test (Figure 2). A single intrathecal dose of the peptidic NTS2-selective JMV-431 (Figure 2A) reversed the cancer-induced allodynia almost completely (anti-allodynic effect of 97% ± 64%, p<0.05; n=9) at the early stage, when compared to the vehicle saline injection. However, an additional dose of JMV-431 at a later stage of pain development did not exert a significant effect (anti-allodynic effect of 40% ± 36%, not significant). However the non-selective agonist JMV-2009 (Figure 2B) induced a significant anti-allodynia on both the early (51% ± 14%, p<0.01; n=9) and late stages of the experiment (33% ± 14%, p<0.05).

**Figure 2.**
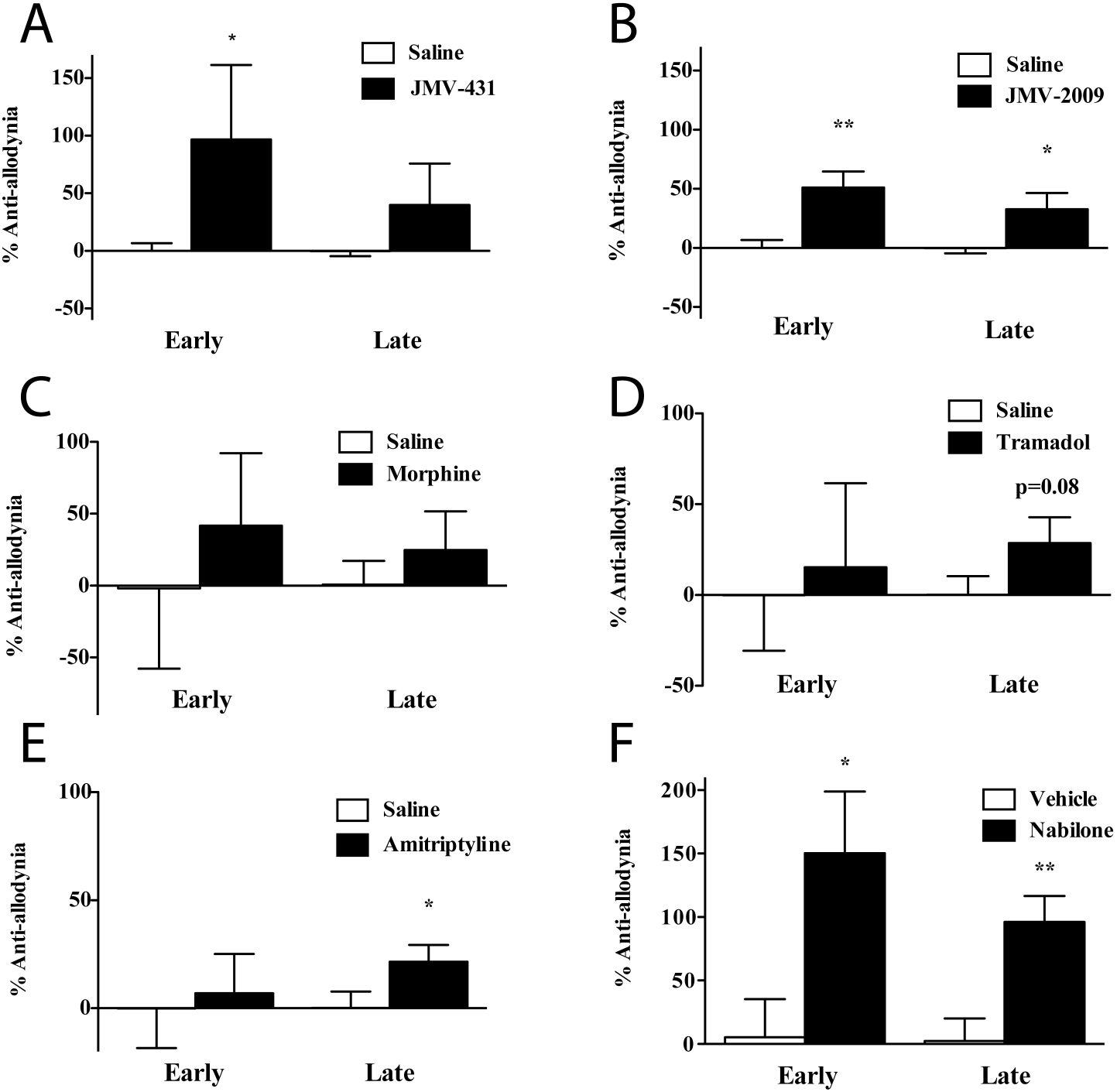
Anti-allodynic effect of six analgesic compounds in BCP. The percentage of anti-allodynic effect is shown for NT agonists A) JMV-431, i.t. 90 µg/kg (NTS2-selective), B) JMV-2009, i.t. 300µg/kg (non-selective) and for other analgesic such as C) Morphine, s.c. 3 mg/kg D) Tramadol, i.p. 15 mg/kg E) Amitriptyline, i.p. 10 mg/kg and F) Nabilone, p.o. 1 mg/kg. All groups are compared to their vehicle group with the same route of administration. *: p ≤ 0.05; **: p ≤ 0.01.

We also used several drugs that are recommended by regulatory authorities and used in practice, at well-documented doses (see table 1)(Dworkin et al., 2010). Subcutaneous morphine (3 mg/kg; Figure 2C) did not significantly reverse tumor-induced allodynia in our model at early (anti-allodynic effect of 42% ± 50%, not significant; n=7) nor late stage (anti-allodynic effect of 25% ± 27%, not significant). The multivalent synthetic analgesic tramadol, a first-line treatment of neuropathic pain (Figure 2D) also had very mild effects on mechanical allodynia. At the early stage of the cancer pain model, no significant effect was observed (anti-allodynic effect of 15% ± 46%, not significant; n=9) compared to the vehicle while a slightly better, but still not significant effect, was observed at late stage (anti-allodynic effect of 29% ± 14%, p=0.08). Similarly, the tricyclic antidepressant amitriptyline, another first-line treatment for neuropathic pain, (Figure 2E) did not show any effect on early pain manifestation (anti-allodynic effect of 7% ± 18%, not significant; n=10). Surprisingly, a second dose administered at an advanced stage significantly reversed mechanical hypersensitivity compared to the vehicle group (anti-allodynic effect of 22% ± 8%, p<0.05). Finally, the cannabinoid nabilone administered orally had a potent significant anti-allodynic effect at both early (150% ± 48%, p<0.05; n=7) and late stage (96% ± 21%, p<0.01).

### Weight bearing modifications following analgesic treatments

Immediately after the von Frey, we performed a DWB test to evaluate the potential of the analgesics on one of the spontaneous components of bone cancer pain (Figure 3). JMV-431 reversed significantly the decrease in weight carried by the ipsilateral paw during the early settlement of pain (Weight recovery of 68% ± 18%, p<0.05) but not during the late stage (Weight recovery of −9% ± 30%, not significant). The JMV-2009 could not induce significant weight recovery at any time point tested with nonsignificant recoveries of 10% and 24% at early and late stages, respectively. Morphine was able to significantly reverse weight shifting to the contralateral paw at the late stage (Weight recovery of 53% ± 18%, p<0.05) of the protocol but not at the early stage (Weight recovery of 30% ± 49%, p<0.05). On the opposite, the tramadol provoked an important, but not statistically significant aggravation of the state of the animal on the DWB (Weight recovery of −117% ± 43%, not significant; p=0.06). The same phenomenon was observed with amitriptyline at an early stage of BCP (Weight recovery of −96% ± 63%, not significant). Nabilone, however induced an important weight recovery during both early (Weight recovery of 101% ± 21%, p<0.05) and late stage (Weight recovery of 60% ± 24%, not significant).

**Figure 3.**
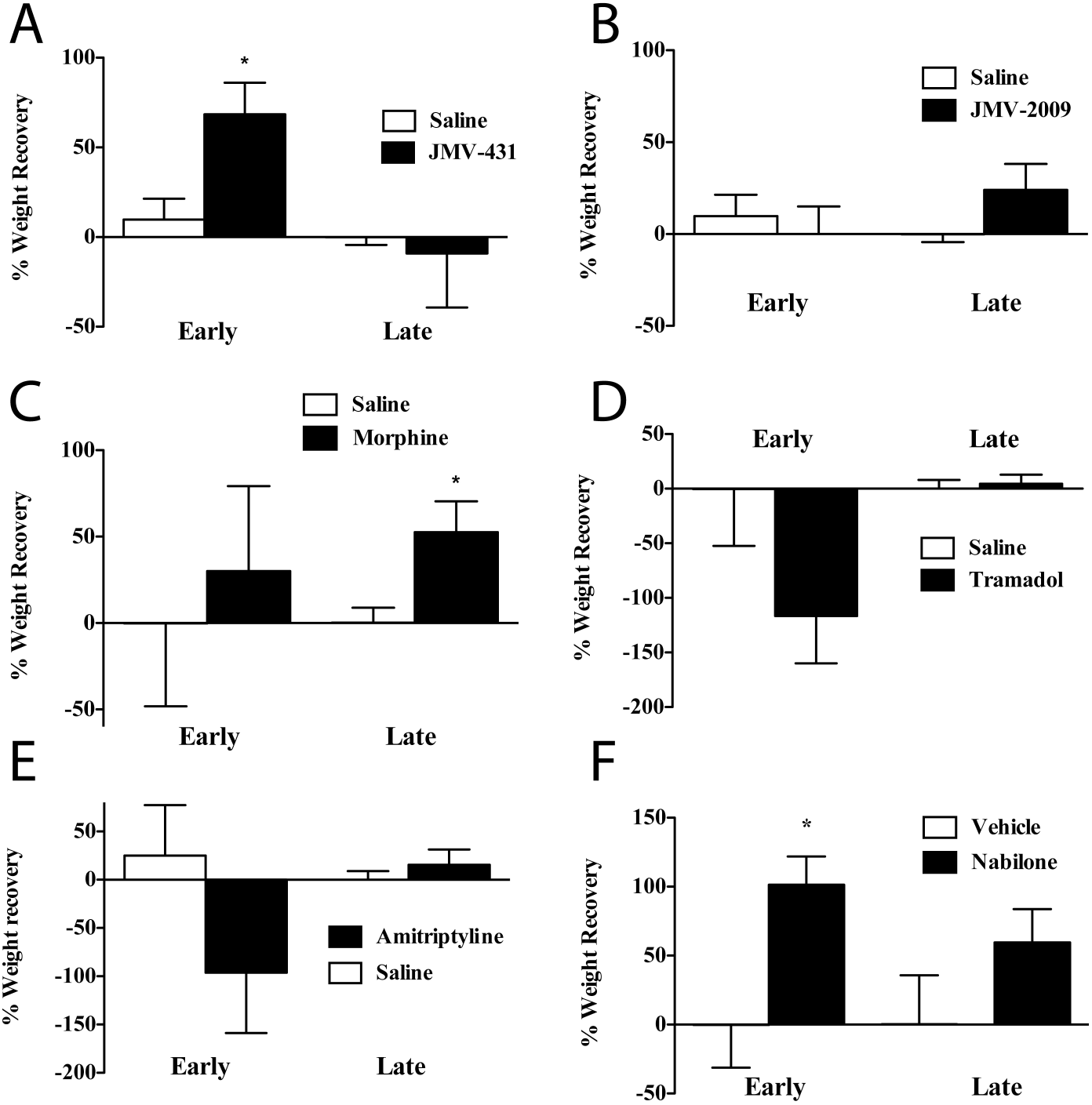
Effect of the analgesics on weight bearing deficits induced by BCP. The percentage of weight recovery effect is shown for NT agonists A) JMV-431, i.t. 90 µg/kg (NTS2-selective), B) JMV-2009, i.t. 300µg/kg (non-selective) and for other analgesic such as C) Morphine, s.c. 3 mg/kg D) Tramadol, i.p. 15 mg/kg E) Amitriptyline, i.p. 10 mg/kg and F) Nabilone, p.o. 1 mg/kg. All groups are compared to their vehicle group with the same route of administration. *: p ≤ 0.05.

### Expression of Neurotensin and its receptors during bone cancer pain progression

We examined the variations of the expression of neurotensinergic system mRNAs at a late stage of BCP (day 18), using real-time quantitative reverse transcriptase PCR (Figure 4). The modifications in expression were assessed for the NT peptide, NTS1 and NTS2 receptors in both L1-L3 lumbar spinal cord (Figure 4A) and lumbar dorsal root ganglia (DRGs) (Figure 4B). In both regions, we did not observe any significant modification in NT and NTS2 expression levels. We observed trends toward an increased expression of NT (139% ± 20% level of expression compared to sham animals, not significant; n=6) and a substantial decrease in the ipsilateral spinal cord for NTS1 mRNA (81% ± 11% level compared to shams, not significant).

**Figure 4.**
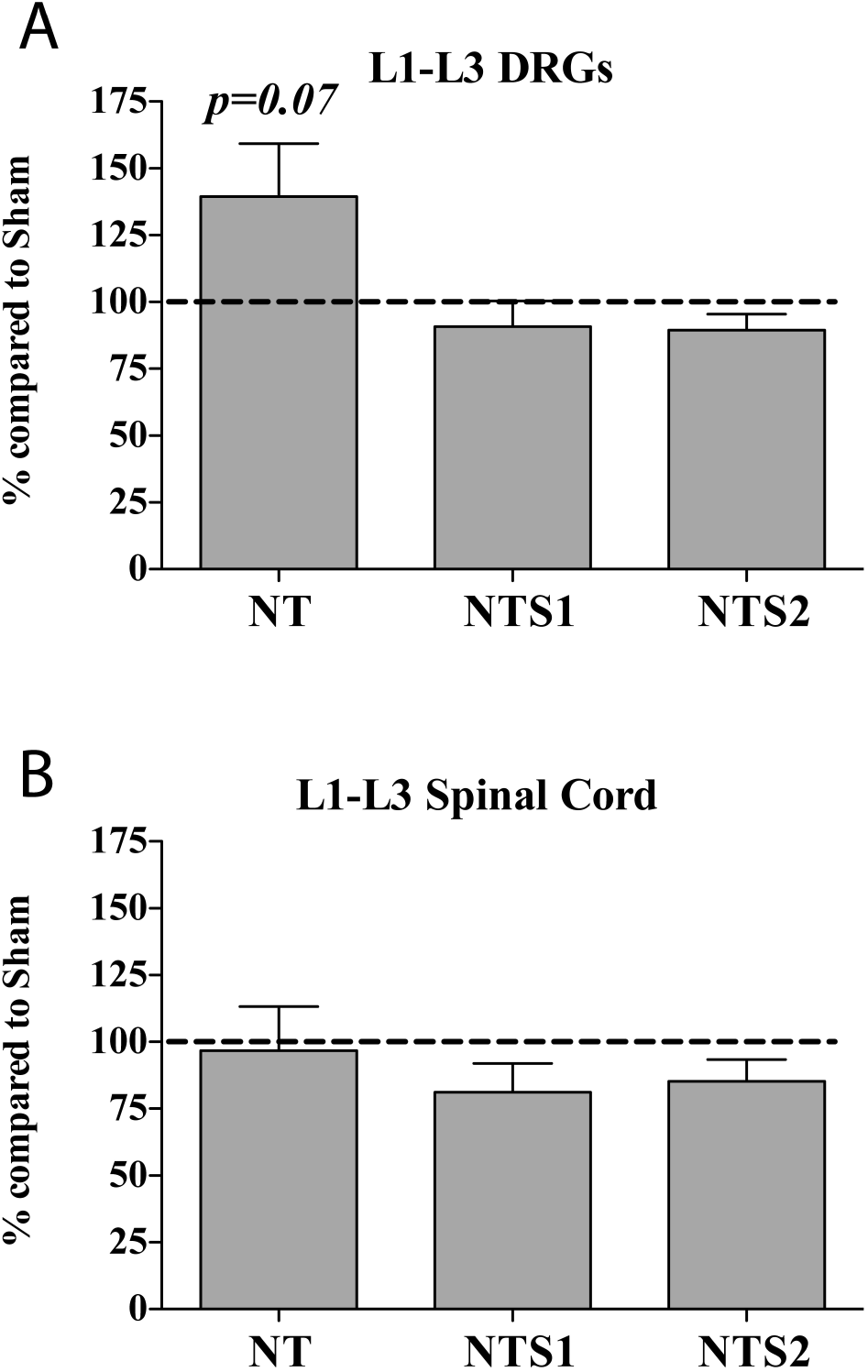
Levels of expression for NT system mRNAs in late BCP context. The expression of NT, NTS1 and NTS2 mRNA was compared between tumor-bearing and sham-operated rat on day 18 post-surgery (late stage). A) In the L1-L3 DRGs, we could not demonstrate a significant variation after BCP onset, despite an interesting increase for NT. B) We observe the same situation in the spinal cord, where NTS1 expression decreased, but not significantly.

## Discussion

In a large proportion of patients, bone cancer pain remains an unsolved puzzle for clinicians. Many analgesic treatments, although satisfactory in other complex pain syndromes, remain useless in the bone tumor context. For instance, morphine and NSAIDs help control background pain, but cannot relieve high intensity breakthrough pain(Mercadante et al., 2009). In many cases, the doses necessary to achieve satisfactory relief are high and patients are under-medicated to minimize intolerable side effects(Mercadante et al., 2007; Nicholson, 2003; Pergolizzi et al., 2008).

The neurotensin system has been under investigation in pain research for 3 decades(Clineschmidt et al., 1979; Dobner, 2006). Our recent work and that of others has demonstrated that neurotensinergic agonists are analgesic leads for treating chronic pain. This study is thus the first one to demonstrate the efficacy of NT agonists in bone cancer pain and supports previous demonstrations of its efficacy in other chronic pain models (Guillemette et al., 2012; Tétreault et al., 2013). However, the advantages of receptor selectivity remain unclear, since non-selective agonists are efficient in certain models and NTS1 or NTS2 selectivity also showed benefits in a few studies, including the present study(Guillemette et al., 2012; Roussy et al., 2008; 2009). Moreover, our group and others are still investigating for compounds that would be available systemically, at least through intravenous injection(Demeule et al., 2014). Meanwhile, we have been testing several compounds to optimize efficiency and minimize side effects. JMV-431 is one of the main candidates and has met success in acute and tonic pain(Doré-Savard et al., 2008; Roussy et al., 2009; Sarret et al., 2005). Its selectivity for NTS2 circumvents typical side effects such as hypothermia and motor deficits, for NTS1 agonists. Our results demonstrate that, in bone cancer pain, JMV-431 is strongly reversing pain at an early stage of the experiment on the von Frey test and, interestingly, in weight bearing paradigm. Postural parameters could be important indicators of analgesic efficiency in animal models of chronic pain(Cobos et al., 2012; Robinson et al., 2012). Indeed, they could be used as a proxy to quality of life in human patient and represent a useful tool to assess the pain relieving potential of a new drug target. These results are in line with previous reports using JMV-431 in the formalin test in mice(Roussy et al., 2009). However, this first attempt at using a non-evoked pain test in chronic pain could pave the way to further developments in the synthesis of BBB-crossing agonists(Demeule et al., 2014; Fanelli et al., 2015; Thomas et al., 2016). Indeed, the JMV-431 can only be injected intrathecally, which limits the possibilities for wide distribution in humans.

In addition to an NTS2-selective peptide, we also tested the novel JMV-2009, a non-selective compound that binds NTS1 and NTS2 with similar affinities. JMV-2009, while showing interesting effects in anti-allodynia at both early and late stages of cancer pain, did not produce any effect on the dynamic weight bearing test. In addition, the peptide did not produce hypothermia despite its analgesic effects. It is an intriguing result raising questions on the factors responsible for weight bearing re-establishment and NTS1-induced hypothermia. First, one could postulate that weight bearing modifications are regulated by NTS2 only and not NTS1. However, JMV-2009 should also be stimulating NTS2 as a non-selective agonist. Moreover, NTS1 agonists showed important effects on neuropathic pain in a chronic constriction injury model(Guillemette et al., 2012). We were thus expecting that a mixed agonist would produce a significant effect on that test, relieving the neuropathic component of cancer pain. However, unpublished results with an NTS1-selective agonist in a similar model in the mouse show the lack of effect of NTS1 stimulation only. It thus appears that NTS2 stimulation alone is better to reverse weight bearing modifications than combined stimulations of NTS1 and NTS2. Our results also support the strong possibility that NTS2 could be an interesting target for neuropathic pain. Such experiments were performed by our laboratory in a recent study(Tetreault et al., 2013).

The expression of NT receptors has been characterized in the spinal cord and dorsal root ganglia in normal conditions(Roussy et al., 2008; Sarret et al., 2005). However, the existence of modulations in their expression in pain context has not been established. In this study, we quantified the mRNA expression for NT peptide and NTS1 or NTS2 receptors in these structures at a late stage of pain settlement. According to our results, it appears that the NT system is weakly regulated in presence of advanced BCP, as it has been observed by others in different chronic pain model(Xiao et al., 2002; Zhang et al., 1996). However, it is not excluded that a time-dependent regulation of expression for some molecules in the spinal cord could take place in BCP. Indeed an early regulation could occur, as it is the case for microglial markers in early neuropathic pain models (Hald et al., 2009). Neurotensinergic system modulations have also be observed in early settlement of neuropathic pain(Vachon et al., 2004; Xiao et al., 2002; Yang et al., 2004).

It is worth mentioning that in both Xiao and Yang studies, only a decrease of NTS2 expression in the spinal cord was observed. It would be interesting to have a more exhaustive investigation of NTS2 expression in time in both mRNA and protein levels.

In this study, we further evaluated the relevance of our bone cancer model in female rats by first testing the efficiency of pain medications that have met variable levels of success in bone cancer pain relief in animal models and humans. As a result, punctuate administrations of morphine, tramadol and amitriptyline succeeded to alleviate pain in a limited number of situations. This has been observed in the mouse, where a dose of 30 mg/kg of morphine was necessary to reach a significant level of anti-allodynia at an advanced stage of tumor development(Luger et al., 2002). Unexpectedly, we observed that morphine treatment at a lower dose was useful in our model only at a later stage of the experiment for weight bearing recovery. This is an interesting observation considering the high level of pain displayed by the animals at that stage; morphine was the only compound able to reverse weight bearing deficits at that stage. Medhurst and colleagues observed the opposite in a study where the same dose reversed mechanical allodynia at an earlier stage of tumor growth(Medhurst et al., 2002). On the other hand, their results in weight bearing, using a static apparatus, are consistent with what we observed. However, the effect of morphine in our study was highly inconsistent from one animal to the other.

On the other hand the tricyclic anti-depressant amitriptyline, used as a first line treatment in neuropathic pain syndromes, was able to reverse mechanical allodynia in a consistent manner at late stage of the disease but it could not reverse (and to some extent worsened) weight bearing modifications. The reason why it was effective for allodynia at that stage compared to an earlier time point is unclear and an accumulation of effect with two consecutive doses cannot be excluded. We also expected tramadol to exert an interesting effect on bone pain, considering its action on both the opioidergic system and on serotonin re-uptake, a frequent treatment in neuropathic pain(Sindrup et al., 2005). However, in the present study, tramadol effects were not significant on both tests and at both time points investigated. The choice of the dose was based on several previous reports (see materials and methods section) and might have been insufficient. Indeed, this study presents the first attempt of using tramadol in a rat model of bone cancer pain and it appears that titration would be necessary. For instance, a dose of 50 mg/kg was necessary to obtain a significant effect in a mouse model(Mouedden and Meert, 2007). It would thus be interesting to use a higher dose in our model, if such a dose can be used in rats.

This study is also the first one to use oral nabilone as a treatment for bone cancer pain in an animal model. This systemically available cannabinoid has been tested in humans for the treatment of neuropathic pain syndromes such as fibromyalgia, multiple sclerosis-and diabetes-induced neuropathy. Our results show that nabilone is one of the two compounds tested that were able to reverse mechanical allodynia at both time points. These results are in accordance with several studies using different cannabinoid compounds(Hald et al., 2008; Hamamoto et al., 2007; Khasabova et al., 2008; Potenzieri et al., 2008). A recent study by Lozano-Ondoua also demonstrated the ability of CB2 agonists to reverse bone cancer pain(Lozano-Ondoua et al., 2010). Moreover, an important effect on the disease was observed by a decrease in bone loss after repeated intra-peritoneal administrations. It would be highly interesting to see if orally-available nabilone could share these same effects on bone loss through the imaging techniques displayed in Figure 1. However a completely different experimental paradigm should be used with daily or bi-daily regimen, as it was shown in the paper of Lozano-Ondoua(Lozano-Ondoua et al., 2010).

It is obvious, in light of our present results and available clinical data, that a single analgesic will not solve the problem of cancer pain. However, publications such as this one, where novel non-evoked pain assessment test are used, could provide insight on effective drug combination to reduce several aspects of bone cancer pain, in addition to complement other studies using non-evoked pain tests such as the catwalk or conditioned place preference(Angeby-Möller et al., 2008; Ding et al., 2005; King et al., 2009). Recent work by Boules and colleagues showed synergy between the effects or a neurotensin agonist and morphine(Boules et al., 2009; 2011). With the results showed in this study, one avenue to consider would be cannabinoids/neurotensin interactions. Indeed, we obtained promising results with Nabilone and with JMV-431, without reaching full relief, especially at late stage. The combination of Nabilone with a neurotensin agonist has the potential to be a powerful tool for scientists and clinicians.

## Conclusion

In this study we characterized the effect of several proven analgesics in our new BCP model. This further characterization will help to assess the potential of new analgesic targets in preclinical setting, in hope to translate to human studies in the future. In that respect, we showed that the neurotensinergic system is an interesting target and that NT agonists exert an analgesic effect that is comparable or superior to common doses of well-proven analgesics. We will continue our quest for better, systemically available agonists and investigate alternate treatment patterns, including combinations with opioids and cannabinoids, in order to improve BCP patients’ quality of life.

## Acknowledgements

The authors would like to thank Martin Lepage, Ph.D.; Luc Tremblay, Ph.D.; Roger Lecomte, Ph.D. and Jean-François Beaudoin, Eng. from the Centre d’imagerie moléculaire de Sherbrooke for their assistance with imaging of the model. We also thank Geneviéve Roussy, Ph.D. and Orlane Ballot for their assistance with data analysis.

